# Dynamics of Brain Lateralization during Chinese Natural Speech Processing under the Influence of Sex Hormones: a 7T fMRI study

**DOI:** 10.1101/2023.11.29.569155

**Authors:** Ruohan Zhang, Shujie Geng, Xiaoqing Zheng, Wanwan Guo, Chun-Yi Zac Lo, Jiaying Zhang, Xiao Chang, Xinran Wu, Jie Zhang, Miao Cao, Jianfeng Feng

## Abstract

Though language is considered unique to humans with left dominant lateralization in the brain, the dynamic nature of the interplay between hemispheres during language processing remains largely unknown. Here, we investigated whole-brain functional dynamic lateralization patterns during Chinese language processing and potential sex disparities using functional MRI data of 20 subjects listening to narrative stories in a 7T MRI scanner. Our findings revealed two distinct dynamic lateralization states, with regions of the language system consistently showing the left lateralization but reversed lateralization for other regions. These two states, characterized by higher-level functioning regions exhibiting left- or right-lateralization, corresponded to the processing of rational and emotional contents, respectively. We observed pronounced inclinations towards the former state in males and the latter state in females, especially during the processing of rational contents. Finally, genetic analyses revealed that the sex differences in lateralization states were potentially influenced by sex hormones.

## Introduction

Languages are distinctively used by humans, and language processing is characterized by intricate, complex, and diverse neural mechanisms. One notable aspect of language processing is brain lateralization, as the language system was traditionally assumed to be situated mainly in the left hemisphere, enhancing efficiency in multitask performance (Esteves, Lopes, Almeida, Sousa, & Leite-Almeida, 2020; Vallortigara, Chiandetti, & Sovrano, 2011) and reducing redundancy in neural organization (Gerrits, Verhelst, & Vingerhoets, 2020; Joliot, Tzourio-Mazoyer, & Mazoyer, 2016; Lazard, Collette, & Perrot, 2012; McAvoy et al., 2015). Nevertheless, some neuroimaging studies have revealed language processing-related regions in the right hemisphere, though they are relatively marginal (Binder et al., 1997; Fedorenko, Hsieh, Nieto-Castañón, Whitfield-Gabrieli, & Kanwisher, 2010; Price, 2012). Furthermore, as a higher-level cognitive function, language processing relies on information coordination across a broad range of cognitive domains, including memory, learning, and cognitive control (Bialystok, Craik, Klein, & Viswanathan, 2004; Doron, Bassett, & Gazzaniga, 2012; Federmeier, Jongman, & Szewczyk, 2020; Fedorenko & Thompson-Schill, 2014; Feng, Chen, Zhu, He, & Wang, 2015; Makuuchi & Friederici, 2013; Novick, Trueswell, & Thompson-Schill, 2010; Peng et al., 2023; Pliatsikas, 2020; Zheng et al., 2020). Most previous studies on language lateralization typically considered language processing a distinct module within the brain. However, the functional lateralization of language processing throughout the brain cortex remains largely unknown.

Moreover, lateralization has traditionally been treated as a static feature of the human brain. However, researchers have recognized that the inherent dynamic coordination of information is a fundamental requirement for optimal brain functioning, spanning both cognitive and behavioural domains (Hiltunen et al., 2014; Kucyi, Tambini, Sadaghiani, Keilholz, & Cohen, 2018; Wirsich, Giraud, & Sadaghiani, 2020). Specifically, functional magnetic resonance imaging (fMRI) studies have revealed that language processing involves highly dynamic interactions across different brain regions (Doron et al., 2012; Fedorenko & Thompson-Schill, 2014; Feng et al., 2015; Peng et al., 2023; Pliatsikas, 2020; Zheng et al., 2020). The processing hierarchy from visual processing to working memory and core language systems during sentence recognition has been revealed with dynamic causal models (Makuuchi & Friederici, 2013). Different dynamic interaction patterns between linguistic and domain-general systems during various cross-language processing operations, such as backwards and forwards translation, have also been reported (Zheng et al., 2020). Notably, by examining the dynamic properties of different brain regions within a predefined language system, researchers found that regions in the left hemisphere have stable processing functions, while those in the right hemisphere have more flexible periphery functions (Chai, Mattar, Blank, Fedorenko, & Bassett, 2016). Nevertheless, our understanding of the dynamic functional lateralization of language processing across the entire cerebral cortex remains limited. In a recent study, Wu et al. (Wu et al., 2022) investigated dynamic lateralization in the brain in the resting state and its association with language functions and cognitive flexibility. This work suggests the potential of exploring dynamic lateralization across the whole brain during language processing.

Furthermore, sex differences have been reported for many cognitive functions, such as theory of mind (Greenberg et al., 2023), visual-spatial processing (Clements et al., 2006), and language processing (Clements et al., 2006; Kansaku, Yamaura, & Kitazawa, 2000; Shaywitz et al., 1995; Xu et al., 2020; Yang et al., 2020). Brain imaging studies have suggested sex differences in brain lateralization during language processing. For instance, a few investigations have indicated that males exhibit stronger left lateralization in the brain than females while performing language tasks (Kimura, 1992; Levy, 1972; Shaywitz et al., 1995), while females exhibit greater bilateral activation in language-related regions than males, including the inferior frontal gyrus, posterior superior temporal gyrus, and fusiform gyrus (Burman, Minas, Bolger, & Booth, 2013; Clements et al., 2006; Kansaku et al., 2000; Xu et al., 2020; Yang et al., 2020). Nevertheless, the impact of sex on dynamic lateralization patterns throughout the brain during language processing remains largely unknown. Additionally, although studies on semantics have unveiled distinct patterns of brain activation across cortical brain regions in response to various language contents (Bi, 2021; Huth, De Heer, Griffiths, Theunissen, & Gallant, 2016), the potential impact on dynamic lateralization, especially in the different sexes, remains unexplored.

To address these issues, we acquired fMRI data using an ultrahigh-field 7T Siemens Terra MR scanner with a cohort of 20 participants (with a mean age of 23.7 ± 1.8 years and 10 females for sex balance) while they performed a Chinese language natural listening task. During the task, all participants listened to two-hour narratives that replicated authentic language to simulate language processing in real-life situations. For the analyses, we employed a data-driven approach that combines a sliding window technique with the global signal-based laterality index to identify recurring whole-brain dynamic lateralization states. Then, we examined the spatial and temporal properties of the states as well as influences of sex and content. Next, we explored the genetic underpinnings of the observed dynamic lateralization states with publicly available postmortem brain-wide gene expression data. Finally, using resting-state fMRI data from the Human Connectome Project (HCP) cohort, we investigated whether the properties of the dynamic lateralization states observed during the Chinese natural listening task are related to inherent lateralization patterns. Figure 1 provides a general schema of the present study.

**Figure 1.**
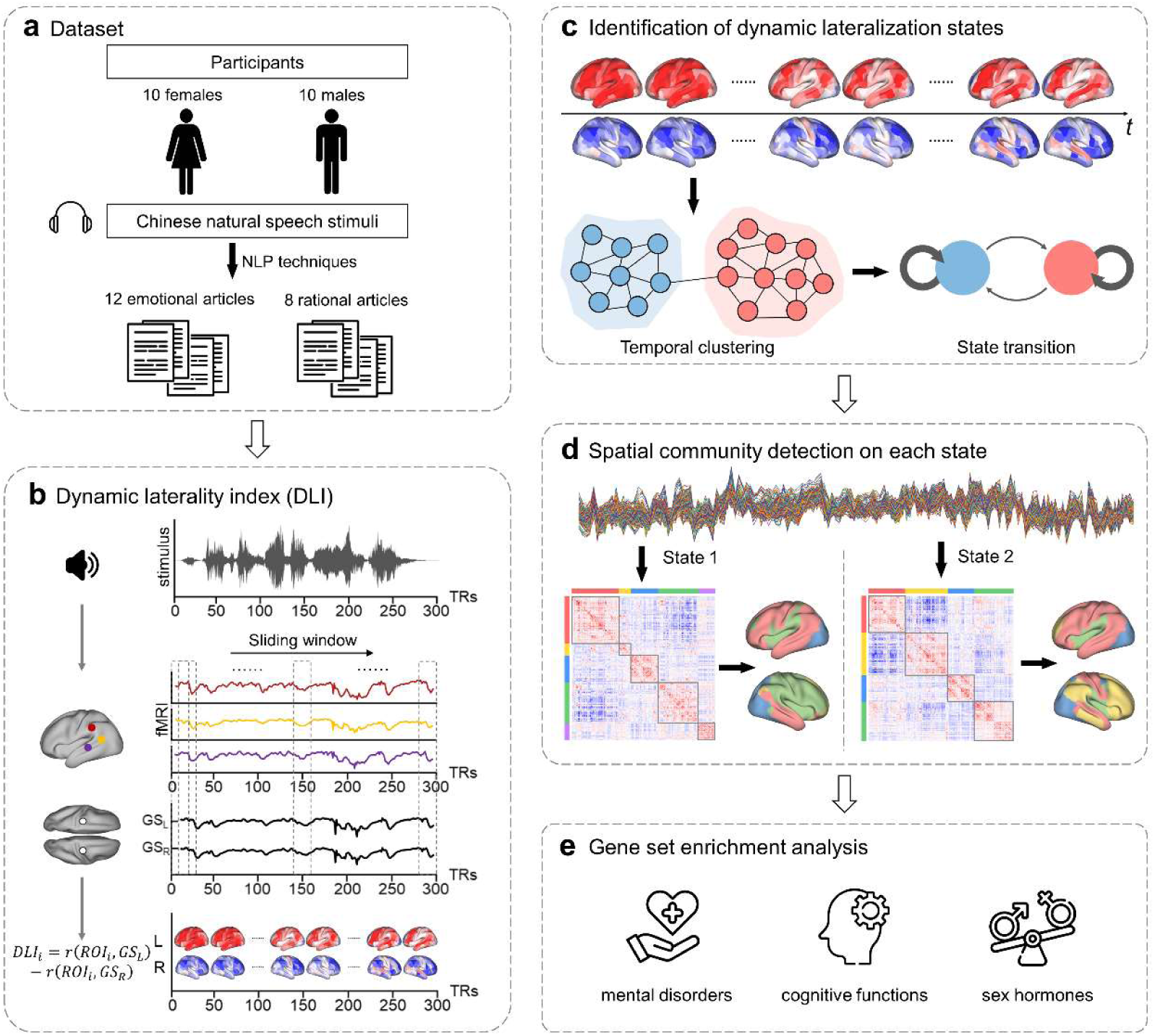
Overview of the study. (a) Task-based fMRI data acquired from a total of 20 participants, including 10 females. Participants were presented with a series of twenty 8- to 12-minute articles, which were categorized into two groups (12 emotional articles and 8 rational articles) using NLP techniques and a hierarchical clustering algorithm. (b) The definition of the DLI. The DLI of *ROI_i_* within the time window *t* is defined as the correlation coefficient (z-transformed) between *GS_L_* and the ROI minus the correlation coefficient between *GS_R_* and the ROI. Using a sliding window approach, we obtained a DLI time series for each ROI. (c) The temporal clustering of the whole-brain lateralization patterns, which identifies recurring dynamic lateralization states. (d) The spatial clustering results of the dynamic lateralization states, which identified spatial communities. (e) The gene set enrichment analysis of the dynamic lateralization states.

## Results

### Categories of the Chinese natural speech stimuli

Participants listened to a set of twenty 8- to 12-minute narratives covering various genres, including fairy tales, fables, novels, essays, news, and scientific essays. We utilized natural language processing techniques and hierarchical clustering analysis approaches to classify the Chinese natural speech stimuli based on their semantic information. The clustering results were visualized using a dendrogram, revealing two distinct article categories (Figure S1a). Then, we performed an assessment to confirm the optimal number of clusters based on the silhouette criterion (Rousseeuw, 1987). This evaluation affirmed the suitability of classifying articles into two categories, as indicated by the highest silhouette value (Figure S1b). Subsequently, we manually labelled one category as articles with an emotional narrative tone (12 articles) and the other as articles with a rational narrative tone (8 articles) (see Table 1 for details). These labels were used to investigate the influence of the type of contents on brain dynamic lateralization patterns.

**Table 1.**
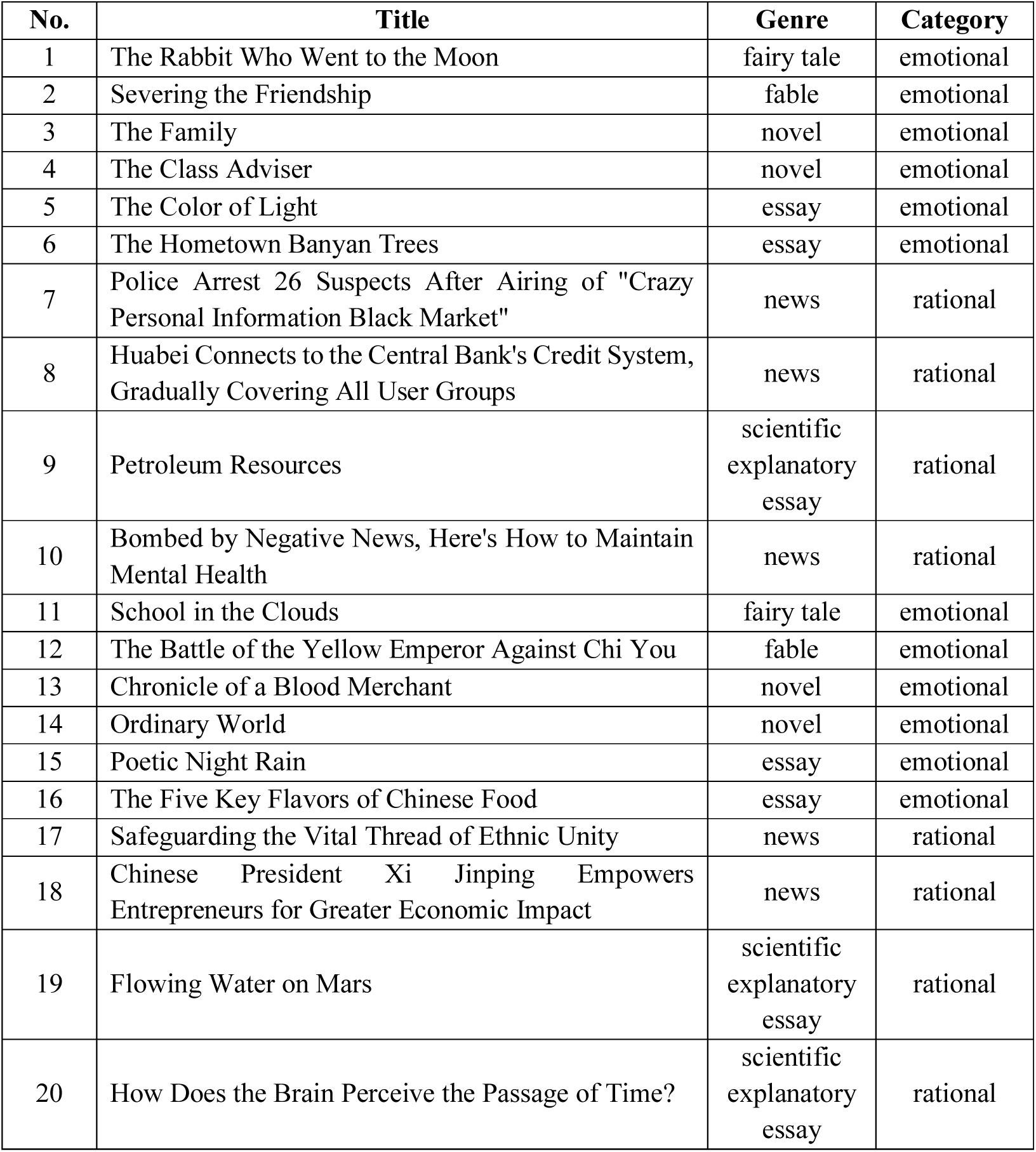
Two categories of the Chinese natural speech stimuli.

| No. | Title | Genre | Category |
| --- | --- | --- | --- |
| 1 | The Rabbit Who Went to the Moon | fairy tale | emotional |
| 2 | Severing the Friendship | fable | emotional |
| 3 | The Family | novel | emotional |
| 4 | The Class Adviser | novel | emotional |
| 5 | The Color of Light | essay | emotional |
| 6 | The Hometown Banyan Trees | essay | emotional |
| 7 | Police Arrest 26 Suspects After Airing of "Crazy Personal Information Black Market" | news | rational |
| 8 | Huabei Connects to the Central Bank's Credit System, Gradually Covering All User Groups | news | rational |
| 9 | Petroleum Resources | scientific explanatory essay | rational |
| 10 | Bombed by Negative News, Here's How to Maintain Mental Health | news | rational |
| 11 | School in the Clouds | fairy tale | emotional |
| 12 | The Battle of the Yellow Emperor Against Chi You | fable | emotional |
| 13 | Chronicle of a Blood Merchant | novel | emotional |
| 14 | Ordinary World | novel | emotional |
| 15 | Poetic Night Rain | essay | emotional |
| 16 | The Five Key Flavors of Chinese Food | essay | emotional |
| 17 | Safeguarding the Vital Thread of Ethnic Unity | news | rational |
| 18 | Chinese President Xi Jinping Empowers Entrepreneurs for Greater Economic Impact | news | rational |
| 19 | Flowing Water on Mars | scientific explanatory essay | rational |
| 20 | How Does the Brain Perceive the Passage of Time? | scientific explanatory essay | rational |

### Two distinct dynamic lateralization states associated with rational and emotional contents processing

We employed an approach combining a sliding window technique and the global signal-based laterality index to calculate the dynamic laterality index (DLI) of the 426 brain regions in the HCPex atlas (Huang, Rolls, Feng, & Lin, 2022). The group-averaged laterality maps across all time windows are shown in Figure 2a. For regions in the left hemisphere, most of them showed left lateralization, except for specific ones in the visual cortex, paracentral lobule, middle cingulate, and posterior cingulate, which showed slight right lateralization. In contrast, nearly all right hemisphere regions displayed right lateralization, except for some auditory association areas (such as the STG area) exhibiting relatively neutral laterality (Figure 2a).

**Figure 2.**
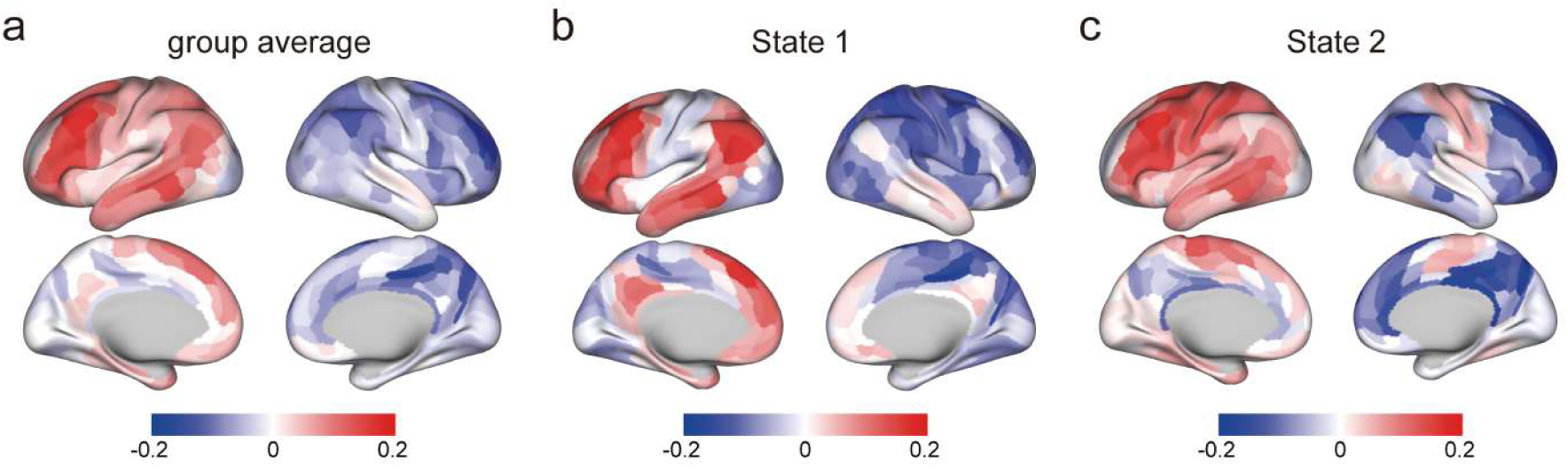
Identification of two dynamic lateralization states. (a) The group-averaged lateralization maps across all time windows, with the majority of brain regions in both hemispheres consistently showing lateralization towards the corresponding side. (b) State 1 of the two dynamic lateralization states, where the left hemisphere exhibited strong left lateralization in regions such as the dorsolateral prefrontal cortex and inferior parietal cortex (such as area 45). (c) State 2 of the two dynamic lateralization states, with the right hemisphere displaying pronounced right lateralization in the dorsolateral prefrontal cortex, posterior cingulate cortex, and inferior parietal cortex. (b) and (c) depicting a distinct reversal in the lateralization pattern of specific brain regions between State 1 and State 2, including the somatosensory cortex, the paracentral lobule, and the middle cingulate cortex. In State 1, these regions in the left hemisphere exhibited right lateralization, whereas in State 2, these same regions within the right hemisphere displayed left lateralization.

By employing a two-stage temporal clustering approach based on the Louvain community detection algorithm (Blondel, Guillaume, Lambiotte, & Lefebvre, 2008), we identified two recurring whole-brain dynamic lateralization states with distinct brain lateralization patterns (Figures 2b and 2c). In both dynamic lateralization states, most brain regions consistently exhibited lateralization towards the hemisphere they belong to. Specifically, in State 1, the left hemisphere regions such as the dorsolateral prefrontal cortex and inferior parietal cortex (such as area 45) exhibited strong left lateralization (Figure 2b). In State 2, right hemisphere regions such as the dorsolateral prefrontal cortex, posterior cingulate cortex, and inferior parietal cortex displayed pronounced right lateralization (Figure 2c). However, we observed a distinct reversal of the typical lateralization pattern in certain brain regions in these two states, such as the somatosensory cortex, the paracentral lobule, and the middle cingulate cortex. In State 1, these regions in the left hemisphere exhibited right lateralization, whereas in State 2, these regions in the right hemisphere displayed left lateralization.

To elucidate the properties of these two states, we investigated their potential differences with respect to text sentiment. We segmented all the audio files into sentences using natural language processing (NLP) techniques and then manually assigned the sentiment score to each sentence. Specifically, sentences with rational contents were labelled 0, and those with emotional contents were labelled 1. After aligning the time windows of the fMRI data with the audio data, we determined the sentiment score for each time window and averaged the scores for each state. The average sentiment score for State 1 was 0.29, which was significantly lower than that for State 2, with an average score of 0.32 (t = -3.3, p = 9.6×10^-4^). According to the predefined criteria for sentiment labelling of the audio data, we classified State 1 as the dynamic lateralization state associated with the processing of rational contents, while State 2 was characterized as the dynamic lateralization state associated with the processing of emotional contents.

### Spatial organization and cognitive functions of the two dynamic lateralization states

To investigate the spatial organization of the two dynamic lateralization states, we conducted a spatial community analysis for each state using the Louvain community detection algorithm (Blondel et al., 2008) (Figure 3a). State 1 exhibited five distinct spatial communities, while State 2 displayed four. To ensure cross-state comparability among these communities, we aligned them by reassigning labels based on their anatomical locations.

**Figure 3.**
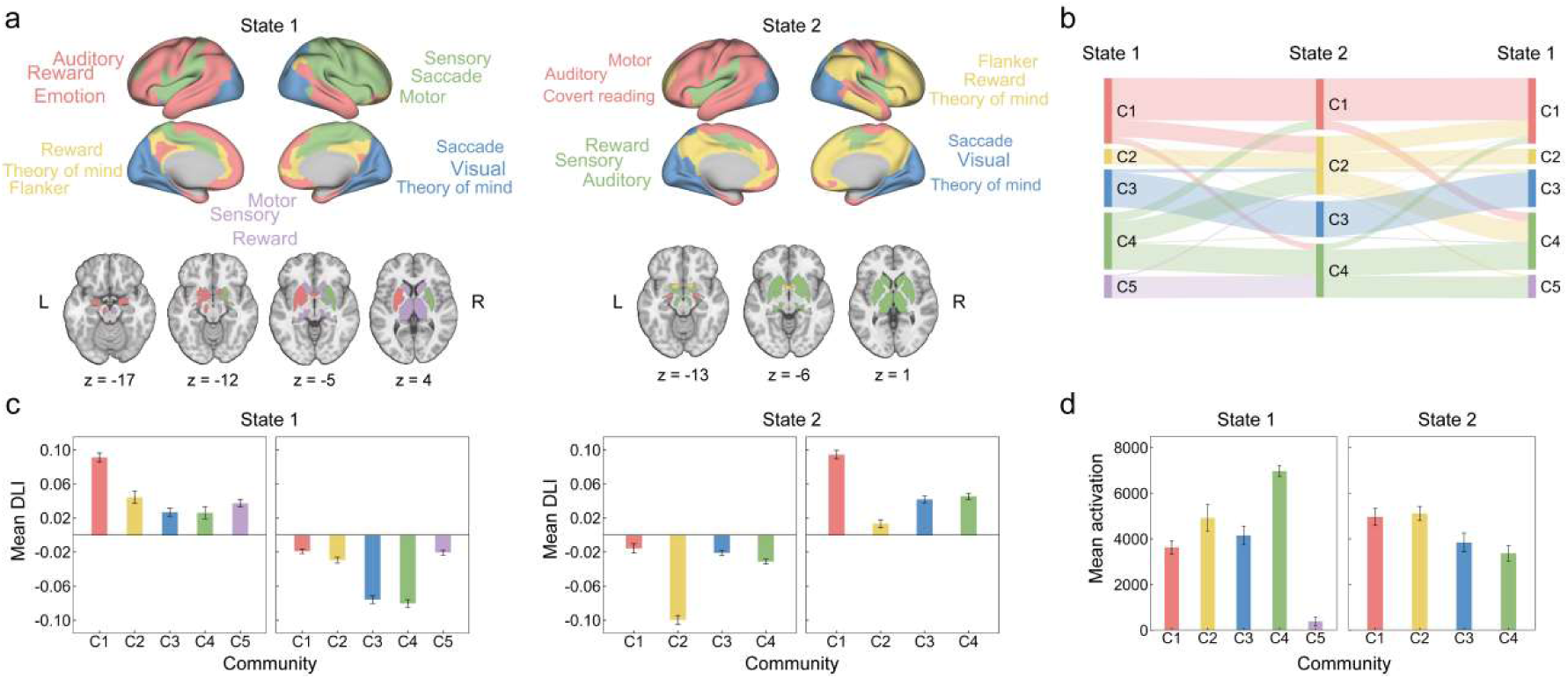
Spatial organization of dynamic lateralization states. (a) The community structures within the two dynamic lateralization states. State 1 exhibits the presence of five distinct spatial communities, and State 2 presents four spatial communities. C1-Language, C2-TOM, C3-Visual, C4-SM, and C5-Subcortical. (b) The segmentation and integration of the communities during the state transitions. (c) The mean DLI values for the communities in these two states. (d) The mean activation values for the communities in these two states. The error bars in (c) and (d) denote the mean ± SEM.

To elucidate the cognitive functions associated with the spatial organization of these two states, we employed a predefined cognitive component template (Yeo et al., 2015) to characterize the cognitive functions associated with each community. In both states, Community 1 (C1), primarily associated with regions for language processing, which was named C1-Language. C2 represented a cluster of regions responsible for higher-level cognitive functions, such as theory of mind, flanker tasks, and reward processing, which was named C2-TOM. C3 was related to lower-level functions such as visual processing and saccades, which was named C3-visual. C4 was associated with sensory and motor processing, which was named C4-SM. C5 was only observed in State 1 and primarily located within subcortical regions involved in sensory and motor processing, which was named C5-Subcortical.

After aligning the communities in the two states, we employed a community transition rate (CTR) matrix to assess how communities are segmented and integrated during state transitions. Overall, C1-Language and C4-SM occupied the large proportions of brain regions in State 1. But in State 2, the proportions of brain regions distributed among the four communities were relatively balanced (Figure 3b). When transitioning from State 1 to State 2, C1-Language and C4-SM in State 1 were segmented and subsequently integrated into other communities in State 2. Specifically, C2-TOM in State 2 included all of C2-TOM, 37% of C4-SM, 26% of C1-Language, 9% of C3-Visual, and 8% of C5-subcortical from State 1. C4-SM in State 2 included half of the brain regions from C4-SM in State 1 and nearly all of C5-Subcortical from State 1. On the other hand, C3-Visual remained relatively stable during the state transition, with most brain regions remaining within their respective communities. Conversely, during the transition from State 2 to State 1, the communities underwent reverse segmentation and integration processes.

The mean DLI values for the communities in these two states are displayed in Figure 3c, showing a significant contrast between the two states. In State 1, C1-Language and C2-TOM clusters showed left-lateralization. While C1-Language was consistently left-lateralization, C2-TOM showed significantly right-lateralization in State 2. Notably, C3-Visual and C4-SM, associated with visual, sensory and motor processing, was predominantly localized to the right hemisphere in State 1 but displayed a slightly left-forward lateralization in State 2. These shifts in hemispheres may be related with the higher prevalence of emotion processing in State 2.

Moreover, we computed the mean activation values for the communities in both states for estimating the functioning involvements (Figure 3d). In State 1, the activation values of the five communities considerably varied, with significant differences in activation observed between every pair of communities after Bonferroni correction (p < 0.05/10). Specifically, in State1, C4-SM exhibited the highest activation, followed by C2-TOM. In State 2, the activation values of the four communities were relatively balanced, yet significant differences in activation were still observed after Bonferroni correction (p < 0.05/6). C1-Language and C2-TOM, which were associated with higher-level functions displayed greater activation values than C3-Visual and C4-SM, which were involved for lower-level functioning.

### Differences in the temporal properties of dynamic lateralization states and influences of sexes and content types

After the exploration of spatial properties, we further examined the disparities in the temporal properties of the two lateralization states. We employed a linear mixed-effects model to calculate the differences of these two states in the occurrence rate, mean dwell time, transition probability, and laterality fluctuation (LF). The influences of both sex and contents (emotional or rational) on these temporal properties were also estimated.

First, we found that the State 2 occurs more than in State 1 with significantly higher occurrence rate (mean of State 1 = 48.2%, mean of State 2 = 51.8%, p = 1.9×10^-3^) (Figure 4a). Moreover, sex and content had considerable impacts on the occurrence rates in both states. Specifically, State 1 happens more in males than females, with significantly differences in occurrence rates (mean of males = 50.2%, mean of females = 46.1%, p = 6.0×10^-3^). Further analysis revealed that this disparity primarily occurred when participants were presented with articles with rational contents (mean of males = 52.0%, mean of females = 46.4%, p = 0.02). Meanwhile, when participants were presented with articles with emotional contents, the sex differences disappeared. Conversely, State 2 occurs more in females than males (mean of males = 49.8%, mean of females = 53.9%, p = 6.0×10^-3^), and this difference was primarily associated with the presentation of rational articles (mean of males = 48.0%, mean of females = 53.6%, p = 0.02). Additionally, no significant difference was observed for only content type in the occurrence rate for either state.

**Figure 4.**
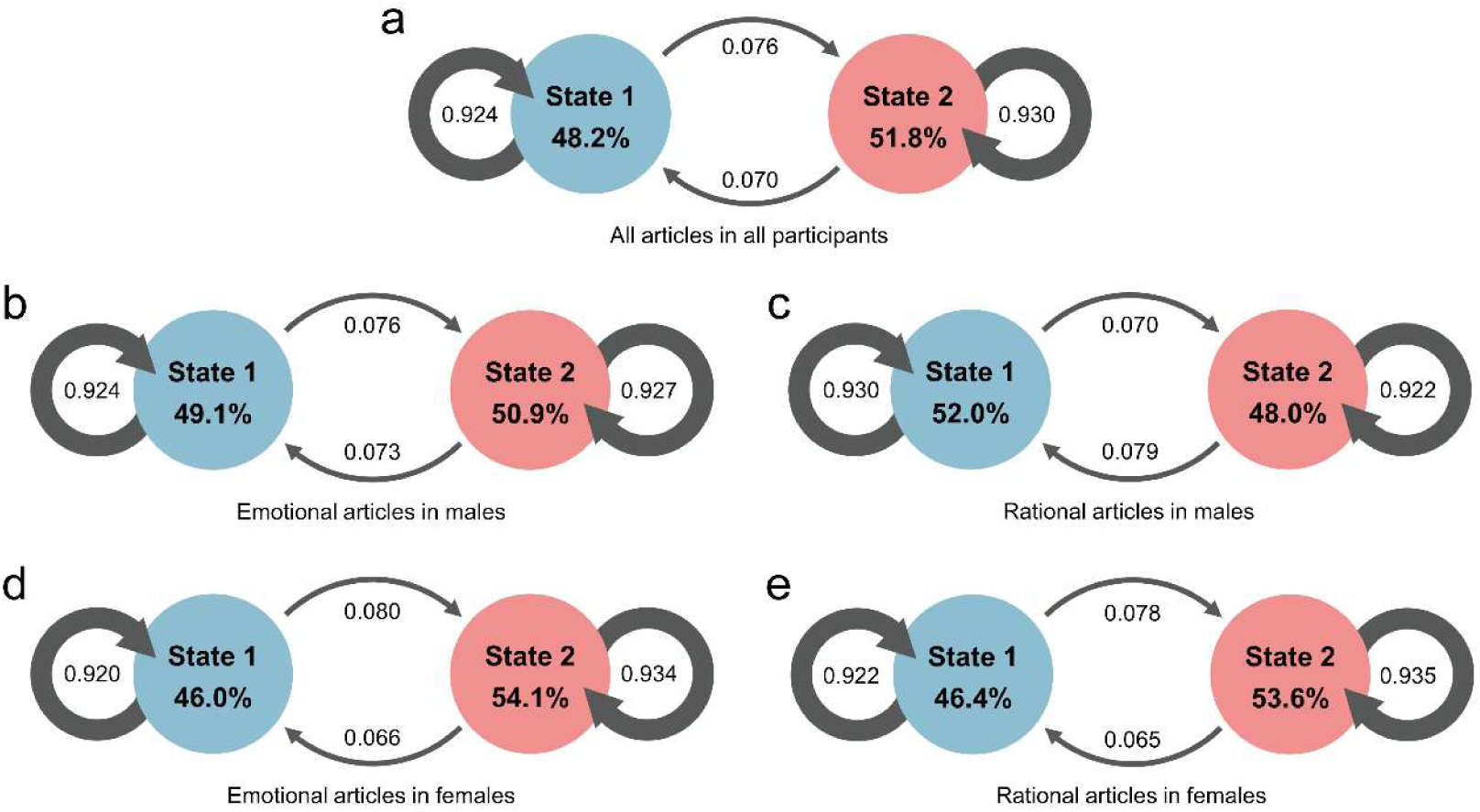
Temporal properties of dynamic lateralization states. (a) The occurrence rates and the mean dwell times of the two states, as well as the transition probabilities between states across all participants and all article sessions. (b) to (e) depicting these temporal properties for different sexes and content types. Linear mixed-effects models were employed to investigate their potential effects on these temporal properties, with age and the number of time windows as covariates in the analysis. The significance of the mixed-effects models was assessed by permutation tests (N = 10000).

Furthermore, the mean dwell time was significantly longer in State 2 than in State 1 (mean of State 1 = 15.6, mean of State 2 = 16.9, p = 0.02) (Figure 4a). This difference was particularly pronounced in females (emotional articles: mean of State 1 = 14.2, mean of State 2 = 17.0, p = 1.0×10^-4^; rational articles: mean of State 1 = 15.5, mean of State 2 = 18.2, p = 8.4×10^-3^) (Figures 4d and 4e). Furthermore, sex differences were observed only in State 1 with males dwelling longer than females (mean of males = 16.4, mean of females = 14.7, p = 0.02). No significant differences in mean dwell time were observed between content types for either state (Figures 4b and 4c).

Additionally, we explored the differences in the transition probability. We found that the probability from States 1 transferred to 2 was significantly higher than that from States 2 to 1 (mean of States 1 to 2 transition = 0.076, mean of States 2 to 1 transition = 0.070, p = 0.01) (Figure 4a). After dividing the population according to sex, this difference was observed only in females during the processing of both emotional articles (mean of States 1 to 2 transition = 0.080, mean of States 2 to 1 transition = 0.066, p = 6.0×10-4) and rational articles (mean of States 1 to 2 transition = 0.078, mean of States 2 to 1 transition = 0.065, p = 0.01) (Figure 4d and 4e). No differences in transition probabilities were observed in males. Furthermore, sex differences were found in the transition probability from States 2 to 1, with the transition probability significantly lower in females than in males (mean transition probability of females = 0.066, mean transition probability of males = 0.075, p = 2.1×10-3). This difference was observed when participants were presented with either emotional (mean transition probability of females = 0.066, mean transition probability of males = 0.073, p = 0.04) or rational articles (mean transition probability of females = 0.065, mean transition probability of males = 0.079, p = 0.01). No sex differences were found in the transition probability from States 1 to 2. Additionally, no significant differences for only content types in transition probabilities were observed.

Significant sex differences in regional laterality fluctuation (LF) were particularly evident during the processing of rational articles with males generally exhibited higher LF values. In State 1, these differences were primarily observed in specific brain regions, including the somatosensory and motor regions, medial prefrontal cortex, posterior cingulate cortex, parahippocampus, insular and amygdala (Figure 5a). In State 2, the brain regions exhibiting significant differences in LF were similar to those in State 1, primarily involving the somatosensory and motor regions, anterior insular cortex, amygdala, dorsolateral prefrontal cortex, and inferior parietal cortex (Figure 5b). Notably, only one region in the insular cortex in the right hemisphere showed significant sex differences in LF during the processing of emotional articles (Figure S2).

**Figure 5.**
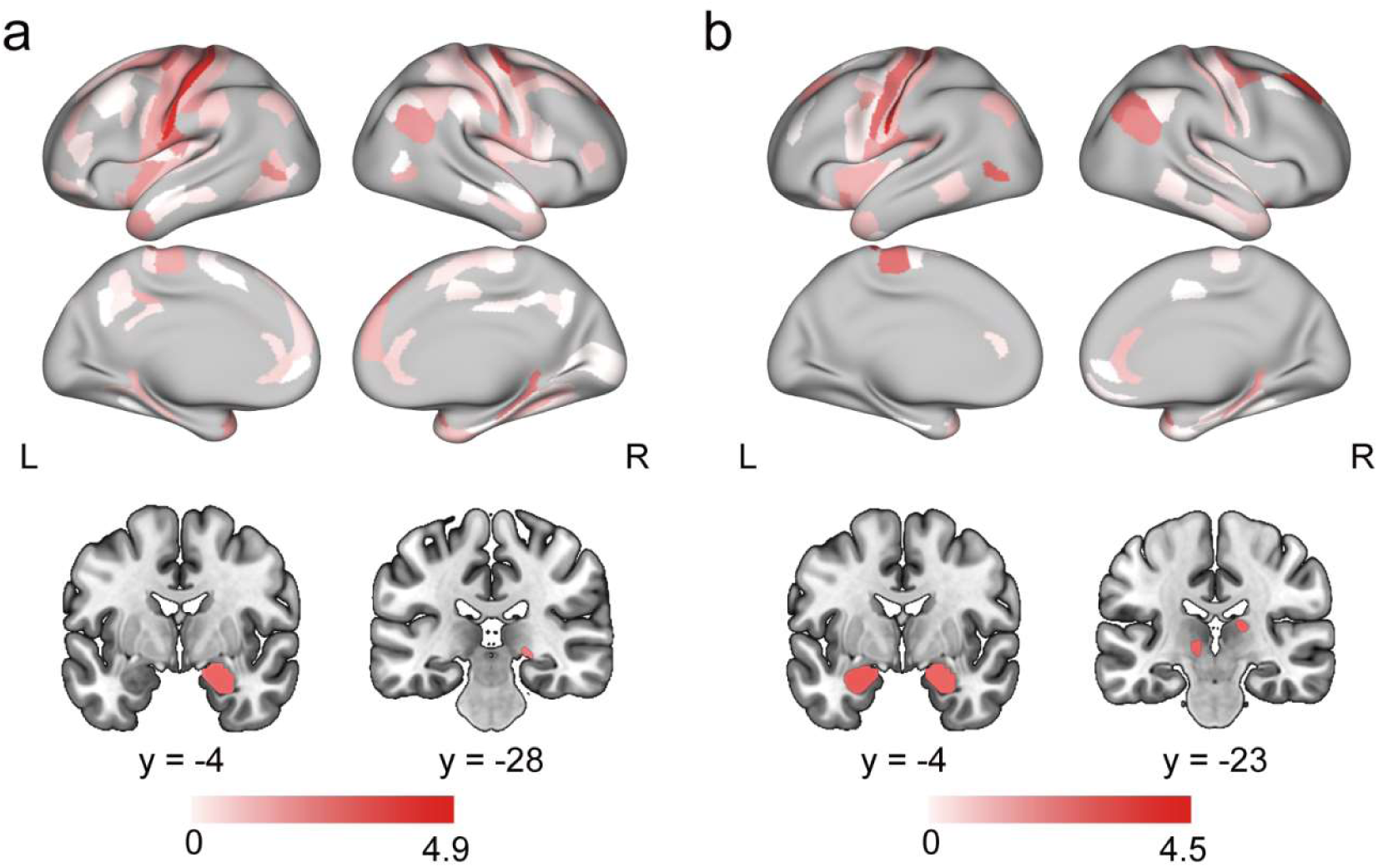
Differences in the laterality fluctuation between sexes (males > females) during the processing of articles with rational contents. (a) and (b) displaying brain regions exhibiting significant differences in laterality fluctuation in State 1 and State 2, respectively. The statistical analyses were conducted using linear mixed-effects models, with age and the number of time windows as covariates. The linear mixed-effects model was fitted according to the LF of each brain region using BH-FDR corrections for multiple comparisons (q value < 0.05).

### Gene set enrichments for dynamic lateralization states

To explore the genetic underpinnings of the two dynamic lateralization states, we conducted a gene set enrichment analysis of the two states. Correlation analyses between gene expression and dynamic lateralization states showed significant associations with State 1 and State 2, and we identified 4,080 and 3,355 genes with significant correlations in States 1 and 2, respectively. These identified genes were associated with specific mental disorders, cognitive functions, and hormones, particularly sex hormones, based on the gene sets from the GWAS Catalog (MacArthur et al., 2017).

More specifically, significant enrichment of genes associated with specific mental disorders, such as neuroticism (q value=7.1×10^-5^), schizophrenia (q value=2.4×10^-4^), and autism spectrum disorder (q value=0.01), was observed in State 1. Furthermore, significant enrichment of genes associated with specific cognitive functions, including general risk tolerance (q value=5.1×10^-7^), cognitive ability (q value=1.2×10^-3^), and educational attainment (q value=0.01), was observed in State 1. Intriguingly, significant enrichment of genes related to hormones, notably male sex hormones, encompassing traits such as chronotype (q value=5.1×10^-7^), sleep duration (q value=4.8×10^-3^), and male-pattern baldness (q value=0.03), was also observed in State 1 (Figure 6a).

**Figure 6.**
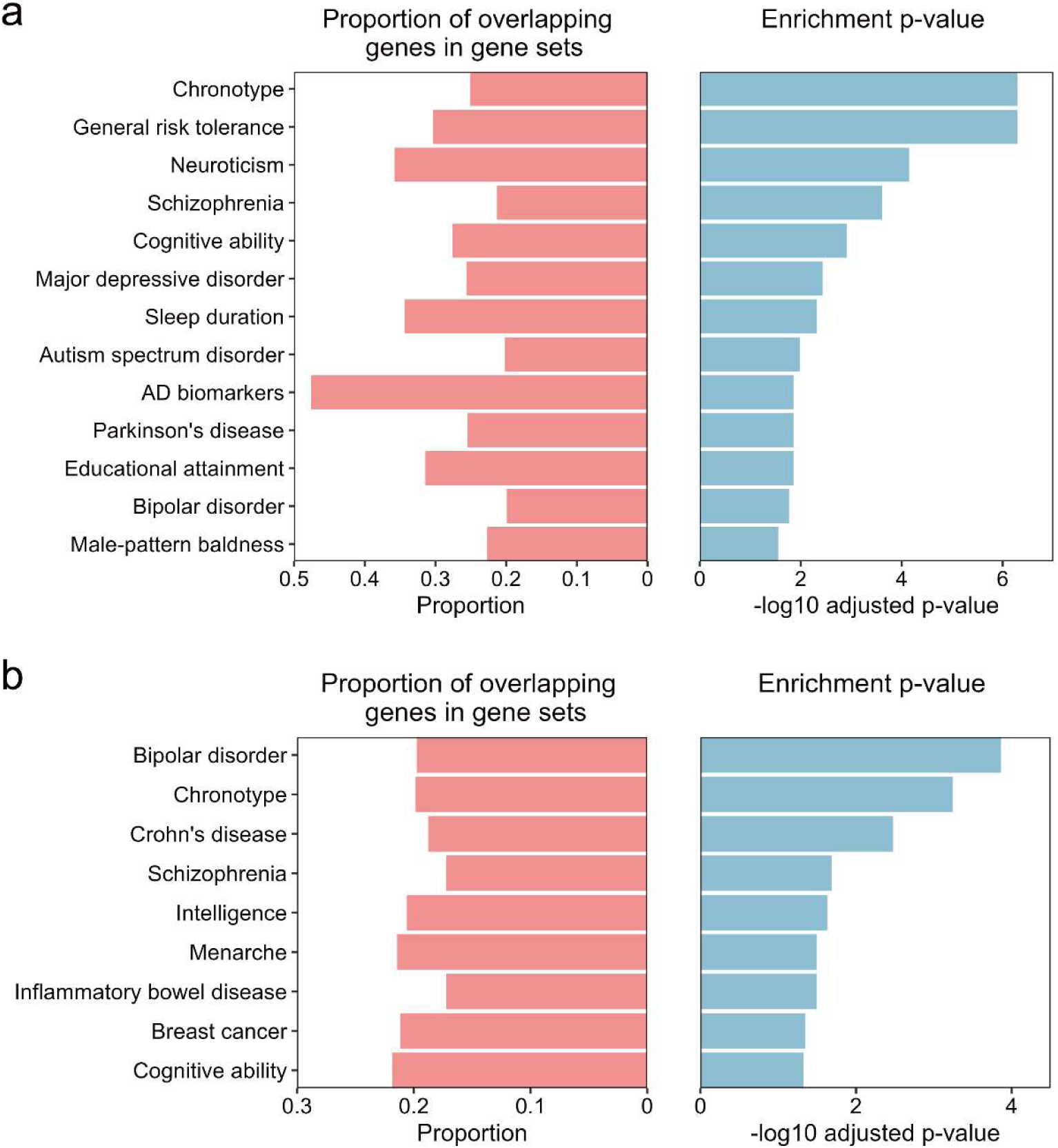
Gene set enrichment analysis for genes associated with dynamic lateralization states. (a) and (b) displaying the biological functions associated with the genes related to the two dynamic lateralization states, as retrieved from the GWAS Catalogue. Multiple test correction was conducted using BH-FDR corrections, with an adjusted p-value cutoff of 0.05 and a minimum of 2 overlapping genes.

In contrast, different gene enrichment patterns were observed in State 2 than in State 1. In addition to genes associated with mental disorders (such as bipolar disorder, q value=1.4×10^-4^) and cognitive functions (such as intelligence, q value=0.02), enrichment of genes associated with inflammatory bowel diseases, specifically Crohn’s disease (q value=3.3×10^-3^), was observed in State 2. Interestingly, significant enrichment of genes associated with female sex hormones, including menarche (q value=0.03) and breast cancer (q value=0.04), was also observed in State 2 (Figure 6b).

### Similarities in dynamic lateralization states between the resting state and Chinese natural speech task

Finally, we would like to investigate whether the dynamic lateralization states identified during the Chinese natural listening task represent inherent lateralization patterns during resting state within the human brain. Here, we conducted an analysis of dynamic brain lateralization using resting-state fMRI data collected from a sample of 991 participants enrolled in the HCP cohort (mean age 28.7 ± 3.7 years, 528 females).

Three distinct dynamic lateralization states were identified during the resting state, as shown in Figure S3. Interestingly, two of these states exhibited significant similarities with the two dynamic lateralization states identified during the processing of Chinese natural speech stimuli (for State 1, r=0.79, p=6.4×10^-79^; for State 2, r=0.79, p=5.3×10^-79^). This implies that the two dynamic lateralization states observed during the processing of Chinese natural speech stimuli reflecting the inherent lateralization patterns within the human brain. These states, characterized by left-lateralized and right-lateralized regions predominantly engaged in higher-level cognitive processes, exhibited heightened activity during the Chinese natural listening task.

Furthermore, the statistical analyses using linear mixed-effects models revealed similar sex differences in temporal properties of these two states. We found that the state, which is similar to State 2 identified during the Chinese natural speech task, occurs significantly more in females than in males (for occurrence rate, t=3.0, p=2.5×10^-3^; for mean dwell time, t=3.3, p=1.2×10^-3^). This finding suggests that sex influences on the temporal properties of the dynamic lateralization states are also inherent, which are similar in both the resting state and during the processing of Chinese natural speech stimuli.

### Validation analysis

To rule out the potential impacts of varying time window lengths, we assessed the reproducibility of the DLI findings using two additional time window lengths, 18 TR and 25 TR. We consistently identified two distinct dynamic lateralization states (Figure S4). Moreover, the two states identified using 18-TR and 25-TR time windows were both remarkably similar to those identified with the 20-TR time window (18 TR: for State 1, r=0.99, p<1.0×10^-15^; for State 2, r=0.99, p<1.0×10^-15^; 25 TR for State 1, r=0.99, p<1.0×10^-15^; for State 2, r=0.99, p<1.0×10^-15^). These results demonstrate the robustness and reliability of the DLI analysis, confirming that our results can be replicated regardless of the chosen time window length.

## Discussion

In this study, we investigated the dynamic nature of whole-brain lateralization during Chinese language processing and explored the potential influences of sex and content (emotional or rational contents), as well as the genetic underpinnings of the observed dynamic lateralization. Remarkably, our findings revealed the existence of two distinct dynamic lateralization states. While the language system was predominantly found in the left hemisphere, intriguingly, we observed converse lateralization patterns in high-level functioning systems. These two states, associated with higher-level functioning systems exhibiting left- or right-lateralization, corresponded to the processing of rational and emotional contents, respectively. More interestingly, males displayed a pronounced inclination for the former state, while females consistently exhibited a preference for the latter state, especially during the processing of rational contents. Furthermore, our investigation revealed genetic evidence that supports these observed sex differences in language processing.

We identified two dynamic states with switched hemispheric specialization, which reflects a core property of human cognition and a marker of efficient functioning. Our study revealed that, brain regions associated with language processing, such as covert and overt reading, were consistently located in the left hemisphere for both dynamic lateralization states. This emphasized the centralized role of left hemisphere for basic language functions, which would benefit the rapid processing (Braga, DiNicola, Becker, & Buckner, 2020; Budisavljevic et al., 2015; Olulade et al., 2020). Moreover, the most notable distinction between these two states is the lateralization patterns of higher-level cognitive systems that assist in language processing in real-life scenarios. Specifically, in State 1, which was strongly associated with the processing of rational contents, brain regions associated with higher-level functions such as reward processing, theory of mind, and emotion showed left lateralization. Conversely, in State 2, which was associated with the processing of emotional contents, the brain regions involved in higher-level cognitive processes such as reward processing, theory of mind, and flanker tasks exhibited right lateralization. This phenomenon showed the coordinated efforts of higher-level cognitive streams between brain hemispheres, which would be superimposed on the basic language processing during language comprehension to enhance cognitive capacity and fluency (Fernandino et al., 2016; Jung-Beeman et al., 2004). It could be inferred that in real life, the realization of efficiently language processing is not limited to one single system, but requires the dynamic and precise cooperation of multi-systems across the whole brain (Labache, Ge, Yeo, & Holmes, 2023). The specialized processing for each hemisphere also echoed with prior studies suggesting that the left hemisphere is predominantly related to mathematical and logical reasoning (Friederici, 2017; Verhelst, Dhollander, Gerrits, & Vingerhoets, 2021; Westerhausen, Kompus, & Hugdahl, 2014), while the right hemisphere is related to emotion processing (Stoyanov, Decheva, Pashalieva, & Nikolova, 2012; H. Watanabe et al., 2015).

Additionally, we observed the noteworthy sex differences in language processing, and these disparities were influenced by the type of contents. Previous research on sex differences in brain lateralization during language processing revealed the tendency for more left-hemisphere engagement in males (Burman et al., 2013; Clements et al., 2006; Packheiser et al., 2020; Xu et al., 2020; Yang et al., 2020). Here, we revealed a more detailed description about the sex differences in language processing. Irrespective of the content type, females consistently exhibited a preference for State 2, characterized by higher-level functions with more right lateralization, as evidenced by significantly higher occurrence rates and a greater inclination to transition towards this state. In contrast, males displayed a pronounced inclination for State 1, characterized by higher-level functions with more left lateralization, particularly during the processing of rational contents. Our findings indicate that during real-life language processing, females tend to employ a perceptual thinking approach, while males tend to employ rational thinking styles. Notably, this intriguing inference provides the neural mechanism explanation for the empathizing-systemizing theory, which suggests that females typically demonstrate greater empathy, while males exhibit a preference for systems-oriented approaches (Baron-Cohen, Richler, Bisarya, Gurunathan, & Wheelwright, 2003; Greenberg et al., 2023; Greenberg, Warrier, Allison, & Baron-Cohen, 2018).

The gene set enrichment analysis based on the two dynamic lateralization states reflect the fundamental questions about the genetic origin and sex differences of language capacities in cognitive neuroscience. The prominent distinction between these states was observed: In State 1, genes associated with male sex hormones, such as male-pattern baldness, were enriched, whereas in State 2, genes associated with female sex hormones, such as menarche and breast cancer, were enriched. This finding showed how the sex hormones would influence the language dynamic lateralization, which are consistent with the findings that males were more inclined towards State 1, while females consistently preferred State 2. Interestingly, prior research has reported the substantial impact of sex hormones on language development (Bowers, Perez-Pouchoulen, Edwards, & McCarthy, 2013; Enard et al., 2002; Haesler et al., 2007; Wermke, Quast, & Hesse, 2018). Additionally, previous studies have indicated the role of biological sex and sex hormones in shaping functional cerebral asymmetries, including language lateralization (Hausmann, 2017; Maney, 2016). Through the comparison with dynamic states during resting-state, our findings elucidate the origin of the two dynamic lateralization states observed during the processing of natural language processing. We inferred that the activation of the dynamic states originates from inherent lateralization patterns in the human brain and heightened during language processing. Additionally, the same sex differences also exist during resting-state, indicating the hemispheric preferences is a sex inherent characteristic. This result provides potential explanations for the sex differences in other cognitive and behavioural domains.

However, it is important to acknowledge some limitations of our study. First, subject to the fact that the scanning time for each subject was very long, this study only included 20 participants. However, each subject was acquiring 20 scans to simulate different language contents that occur in daily life, which would increase the effective sample size. Additionally, we made efforts to achieve sex balance by recruiting 10 female participants. Future studies with larger and more diverse samples would validate our findings. Another limitation is that the individuals in our study were all young people with high-level education backgrounds (mean age 23.7 ± 1.8 years). It would be valuable to investigate the dynamic lateralization states during Chinese language processing in participants of different ages, including children and older adults. This would allow us to explore potential age-related changes in lateralization patterns and their implications for language processing. Furthermore, our analyses were specifically based on the Chinese language. It would be interesting and important to investigate dynamic lateralization states during English language or other language processing to examine the generalizability of our findings across different linguistic contents.

In conclusion, our study provides valuable insights into the dynamic nature of brain lateralization during Chinese language processing. The identification of two distinct lateralization states improves our understanding of interhemispheric coordination during language processing, emphasizing the dynamic shift in higher-level cognitive functions between left and right lateralization patterns. Furthermore, the associations of these lateralization states with sex differences and the processed contents (emotional or rational contents) offer insights into sex-specific patterns in language processing. Moreover, our genetic-level analyses suggest that the observed sex differences in language lateralization may be influenced by sex hormones. Future research should focus on replicating and expanding our findings utilizing larger and more diverse populations, including individuals with different ages and/or neurodevelopmental disorder, as well as investigating the cognitive significance of the observed dynamic lateralization states. As such, understanding the interactions linking the biological underpinnings of dynamic lateralization, cognition and development would have significant implications for both cognitive neuroscience and developmental biology.

## Materials and methods

### Participants and task-based fMRI dataset

Fully informed consent was obtained from all participants by Fudan University, and the research procedures and ethical guidelines were approved by the Fudan University institutional review board. The task-based fMRI data were collected from a total of 20 participants (mean age 23.7 ± 1.8 years, 10 females). All participants were healthy and had normal hearing. The participants were postgraduate students with the same educational level. The task-based fMRI data were acquired using an ultrahigh-field 7T Siemens Terra MR scanner at the Zhangjiang International Brain Imaging Centre (ZIC) of Fudan University, Shanghai, China. A 32-channel Siemens volume coil (Siemens Healthcare) was used for data acquisition. Functional scans were obtained using an echo planar imaging (EPI) sequence with the following parameters: repetition time (TR) = 1,500 ms, echo time (TE) = 25 ms, flip angle = 65°, voxel size = 1.5×1.5×1.5 mm, matrix size = 128×128, field of view (FOV) = 1,920×1,920 mm, slice thickness = 1.5 mm, and number of slices = 96. The T1-weighted anatomical data were collected on a 3T scanner with the following parameters: TR = 2,500 ms, TE = 2.22 ms, flip angle = 8°, matrix size = 300×320, FOV = 240×256 mm, slice thickness = 0.8 mm, and number of slices = 208.

### Data preprocessing

Image preprocessing was performed using Statistical Parametric Mapping-12 (SPM12; http://www.fil.ion.ucl.ac.uk/spm). To ensure T1 equilibrium, several volumes were not recorded prior to the trigger. The volumes were temporally realigned to the middle EPI volume and spatially realigned to correct for head movement. The structural images of each participant were registered to the mean EPI image, segmented, and normalized to Montreal Neurologic Institute (MNI) space. The realigned EPI volumes were then normalized to MNI space using deformation field parameters obtained from the structural image normalization. Finally, the normalized EPI volumes were spatially smoothed using a 6×6×6 mm Gaussian kernel and high-pass filtered to reduce noise. After preprocessing, the cortical grey matter was parcellated into 426 regions, including 360 cortical areas and 66 subcortical areas, using the HCPex atlas (Huang et al., 2022). The HCPex atlas is a modified and extended version of the surface-based Human Connectome Project-MultiModal Parcellation atlas of human cortical areas (HCP-MMP v1.0) (Glasser et al., 2016). This atlas provides a detailed and multimodal parcellation of brain regions, enabling the examination of dynamic lateralization in different brain regions. For each participant, time series were extracted by calculating the mean signal of all voxels within each of the 426 brain regions.

### Chinese natural speech stimuli

In the task, a set of twenty 8- to 12-minute articles were utilized as the Chinese natural speech stimuli. These articles covered a wide range of genres, including fairy tales, fables, novels, essays, news, and scientific essays. A detailed summary of each article is presented in Table S1. Each article was presented during a separate fMRI scanning session; thus, a total of 20 sessions were conducted for each participant. The duration of each scan was customized to match the length of the corresponding article and included a period of silence lasting 2 seconds both before and after the article was presented. The 20 fMRI scans were conducted on different days, with a total of approximately 162 minutes of scanning time. The articles were played to participants using Sensimetrics S14 in-ear piezoelectric headphones.

### Classification of Chinese natural speech stimuli

Data-driven techniques incorporating natural language processing techniques and the hierarchical clustering algorithm were employed to classify the Chinese natural speech stimuli. The NLP techniques applied in this analysis included the "Jieba" Chinese text segmentation system and text vectorization. First, Chinese word segmentation was performed using the *Jieba* word segmentation tool (https://github.com/fxsjy/jieba/). This tool divided continuous sentences into words. Next, each word was represented as a 300-dimensional numerical vector based on pretrained Chinese word vectors (Li et al., 2018). These vectors were trained with rich content features (word, ngram, and character) and large corpora, including Baidu Encyclopedia, Wikipedia_zh, Wikipedia_zh, Sogou News, Financial News, Zhihu_QA, Weibo, and Literature. This training produced accurate word representations in a high-dimensional semantic space. Subsequently, each article was represented as the term frequency-inverse document frequency (TF-IDF) weighted sum of the embedding vectors corresponding to each word. The TF-IDF is a statistical method that calculates the significance of a token or word to a document within a set of documents (Hakim, Erwin, Eng, Galinium, & Muliady, 2014). The TF-IDF score combines the term frequency (TF) and inverse document frequency (IDF). The TF, tf(*t*, *d*), measures how many times a token or word appears in a document:

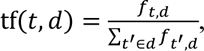

where *f_t,d_* is the number of times word *t* occurs in document *d*, and the sum of *f_t_*!*_,d_*

represents the total number of words in document *d*.

The IDF measures how frequent or rare a token is within the entire document set:

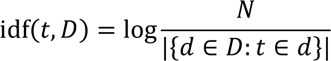

where *N* is the total number of documents and |{*d* ∈ *D*: *t* ∈ *d*}| denotes the number of documents in which the word appears.

By multiplying these two metrics, a TF-IDF score is obtained for each word in a document:

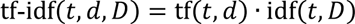

Finally, the hierarchical clustering algorithm (Nielsen, 2016) was applied to the set of 20 vectorized articles. In this algorithm, the Euclidean distance and inverse squared distance (i.e., minimum variance algorithm) are utilized to calculate the distances between clusters. The clustering results were visualized using a dendrogram. The optimal number of clusters was determined by considering both the structure of the dendrogram and the silhouette criterion (Rousseeuw, 1987). The clustering labels obtained with this classification algorithm were then used to investigate differences in dynamic lateralization when processing different types of Chinese natural speech stimuli.

### Dynamic laterality index

To assess the dynamic brain lateralization during the processing of Chinese natural speech stimuli, we utilized an approach proposed by Wu et al. (Wu et al., 2022) for measuring the synchronization changes of brain regions in the left or right hemisphere while performing cognitive functions over time. We used a sliding window method with the window length set as 20 TR and the step set as 1 TR and calculated the dynamic laterality index for each region. For each sliding window, the DLI is defined as:

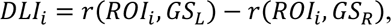

where *ROI_i_* denotes the BOLD time series of ROI *i*, and *GS_L_* and *GS_R_* denote the mean global signal, i.e., the BOLD time series averaged across all voxels, of all regions within the left and the right hemispheres. The Pearson correlation coefficient *r* is Fisher-z-transformed. To test the reproducibility of the DLI, we also employed two other window lengths to evaluate the impact of the window length on the DLI results.

To quantify the magnitude of the time-varying laterality, we measured the LF, which is defined as the standard deviation of the laterality time series (Wu et al., 2022), of each ROI for every participant.

### Identification of dynamic lateralization states

To identify recurring whole-brain dynamic lateralization states, a two-stage temporal clustering approach was employed. In the first stage, we focused on individual-level analysis. For each participant, a temporal similarity matrix was computed across the total 6,114 time windows of the 20 sessions. The temporal similarity was measured as the Pearson correlation between each pair of whole-brain dynamic lateralization states in the different time windows. Then, we applied the Louvain community detection algorithm (Blondel et al., 2008) to the temporal similarity matrix of the 6,114 dynamic lateralization time windows for each participant to generate clusters. The *γ* parameter was set to 1 in the community detection process, and the algorithm was stopped when the modularity Q no longer improved (i.e., improvement < 1.0×10^-100^), ensuring the best cluster partitioning. Each cluster represented a dynamic lateralization state at the individual level, and the state centroids (i.e., the mean of whole-brain dynamic lateralization state within a cluster) were calculated.

In the second stage, we focused on group-level analysis. To obtain group-level recurring dynamic lateralization states, a temporal similarity matrix was computed using the state centroids from all participants. The Louvain community detection algorithm was applied to analyse this matrix. Based on the group-level clustering results, all the dynamic lateralization time windows of all participants were reclassified accordingly.

### Mapping text sentiment to dynamic lateralization states

#### Audio text segmentation and sentiment labelling

The audio data were segmented into sentences using NLP techniques. Regular expression operations were employed to identify sentence boundaries in the audio transcriptions, followed by manual correction. A group of four participants (two females and two males) was recruited to assign sentiment labels to the sentences. These participants were evenly distributed between sexes and were not involved in the task-based fMRI scanning. They rated the sentiment of each sentence, distinguishing between emotional (marked as 1) and rational (marked as 0) sentences. The sentiment ratings provided by the four participants were then averaged to obtain a sentiment score for each sentence. Moreover, we determined the sentiment score of each individual Chinese character within the sentences according to the score of that sentence.

#### Mapping audio texts to functional brain imaging windows

The time windows defined in the task-based fMRI data and the corresponding sections of the audio data were aligned by synchronizing the timestamps of the fMRI data with the timestamps associated with each sentence in the audio data. This temporal alignment enabled precise correspondence between the linguistic information and the neural activity captured in each window.

#### Calculation of sentiment scores for dynamic lateralization states

We collected the Chinese characters in the audio data for each time window in the task-based fMRI data. Subsequently, the sentiment score for each time window was computed by averaging the scores of all Chinese characters within the window, which served as a measure of the emotion of the contents conveyed by the audio data during that particular window. Moreover, to minimize the influence of individual differences on the lateralization state, we determined the lateralization state of each time window by measuring the mode across the 20 participants. This approach established a relationship between the content emotions (emotional or rational) in the audio texts and the corresponding dynamic lateralization states.

### Spatial properties of dynamic lateralization states

#### Community structure of each state

To investigate the spatial organization of the dynamic lateralization states, we first extracted the lateralization time series for each state. The lateralization time series for each state were obtained by concatenating the data from the 20 scanning sessions of the 20 participants. Subsequently, we calculated a spatial similarity matrix (426×426) based on the lateralization time series across the entire brain for each state. The similarity between regions was estimated using the Pearson correlation coefficient, and only relationships with positive weights were retained. Next, we employed the Louvain community detection algorithm (Blondel et al., 2008) to determine the community structure of each dynamic lateralization state based on the spatial similarity matrix. During the community detection process, we set the parameter *γ* to 1, and the algorithm iterations were stopped the modularity Q no longer improved (i.e., improvement <1.0×10^-100^) to obtain the best community partitioning results. To ensure comparability among communities across different dynamic lateralization states, we aligned the communities by relabelling them according to their anatomical locations.

Additionally, we calculated the mean DLI and mean activation values of the communities for each state. Because the language listening task was a continuous process without contrasting tasks, we set the activation threshold as the mean BOLD signal value across the whole brain at each time point. The activation value for each brain region was obtained by thresholding the BOLD signal values.

#### Community segmentation and integration during state transitions

To quantify the process of community change during state transitions, we introduced a community transition rate (CTR) matrix that captures the segmentation and integration of communities during state transitions. Assuming a state space S, the CTR matrix from state *i* with *M* communities to state *j* with *N* communities is defined as:

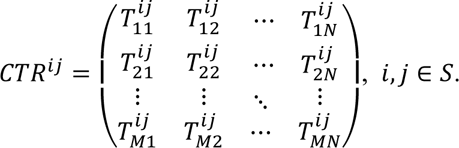

Here, 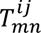 represents the (*m*, *n*) element of *CTR^ij^*, given by:

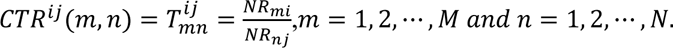

where *NR_mi_* represents the number of brain regions in the *m*th community in state *i*, and *NR_nj_* represents the number of brain regions in the *n*th community in state *j*. Since the matrix elements represent transition rates, they must satisfy:

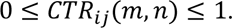

Furthermore, as the brain regions in a community in one state must be allocated to one of the available communities in another state, the transition matrix must satisfy:

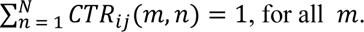

### Temporal properties of dynamic lateralization states

#### Occurrence rate and mean dwell time

The temporal properties of the dynamic lateralization states were assessed using the occurrence rate and mean dwell time (Ryali et al., 2016; Zhou, Zhang, Feng, & Lo, 2019). The occurrence rate represents the proportion of time windows spent in each dynamic lateralization state, measured as a percentage. The mean dwell time indicates the duration of time that a participant remains in a specific dynamic lateralization state, calculated by averaging the number of consecutive windows spent in that state before transitioning to another state.

#### Transition probability

To characterize the transitions between brain states, we estimated the transition probability matrix using the dynamic lateralization state time series for each participant and scanning session. Assuming a state space S with N brain states, the estimated transition probability matrix for the dynamic lateralization state series is denoted as *P*(*N* × *N*). The element (*i*, *j*) in matrix *P* is calculated as:

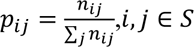

where *n_ij_* denotes the number of times state *i* transitions to state *j*.

### Statistical analysis

We employed linear mixed-effects models (Pinheiro & Bates, 1996) to investigate the potential effects of the temporal properties of the dynamic lateralization states, with age and the number of time windows as covariates in the analysis.

#### Differences in the temporal properties between states

We compared the differences in the occurrence rate among different states using the following model:

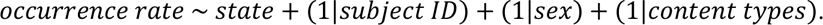

This model included fixed effects for brain lateralization states and random effects for participants, sex, and content types since the task employed a within-subject design. To assess the significance of the explanatory variable in the mixed-effects model, we performed N = 10000 permutations of the response variable. Additionally, we evaluated the differences in the mean dwell time among different states using the same model.

#### Differences in the transition probabilities between transition directions

We compared the differences in the transition probability between different the two transition directions (i.e., States 1 to 2 or States 2 to 1) using the following model:

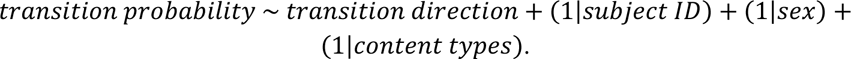

A permutation test (N = 10000) was performed to determine the significance of the mixed-effects model.

#### Differences in the temporal properties between sexes or content types

We compared the differences in the occurrence rate between different conditions (sexes or content types) using the following model:

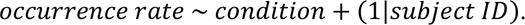

We performed a permutation test (N = 10000) to assess the significance of the mixed-effects model. Similarly, the differences in the mean dwell time (or transition probability) between sexes or content types were assessed using the same approach.

#### Differences in the laterality fluctuation between sexes or content types

We compared the differences in the laterality fluctuation between different conditions (sexes or content types) using the following model:

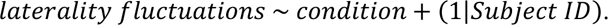

The linear mixed-effects model was fitted for the LF value of each brain region. P-values were corrected for multiple comparisons using Benjamini-Hochberg false discovery rate (BH-FDR) corrections (q value < 0.05) (Benjamini & Hochberg, 1995).

### Gene expression analysis of dynamic lateralization states

#### Gene expression data

Regional microarray expression data were obtained from six postmortem brains (one female, ages 24.0∼57.0 years, mean age 42.50±13.38 years) provided by the Allen Human Brain Atlas (AHBA, https://human.brain-map.org) (M. J. Hawrylycz et al., 2012). The data were processed using the abagen toolbox (version 0.1.3; https://github.com/rmarkello/abagen) and the HCPex atlas, a 426-region volumetric atlas in MNI space.

First, the microarray probes were annotated using the data provided by Arnatkevičiūtė et al. (Arnatkevičiūtė, Fulcher, & Fornito, 2019). The probes that did not match a valid Entrez ID were discarded. Next, the probes were filtered based on their expression intensity relative to background noise (Quackenbush, 2002). Probes with intensities lower than the background in >=50.00% of samples across donors were discarded. In cases where multiple probes matched the expression of the same gene, we selected the probe with the most consistent pattern according to regional variations across donors (i.e., differential stability) (M. Hawrylycz et al., 2015). The differential stability was calculated using the formula:

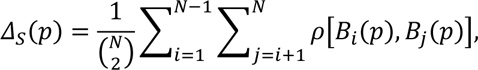

where *ρ* is the Spearman’s rank correlation of the expression of a single probe, *p*, across regions in two donors *B_i_* and *B_j_*, and *N* is the total number of donors. The regions corresponded to the structural designations provided in the ontology from the AHBA.

The MNI coordinates of the tissue samples were updated by performing nonlinear registration using advanced normalization tools (ANTs; https://github.com/chrisfilo/alleninf). Samples were assigned to brain regions in the provided atlas if their MNI coordinates were within 2 mm of a given parcel. To minimize assignment errors, sample-to-region matching was constrained by hemisphere and gross structural divisions (i.e., cortex, subcortex/brainstem, and cerebellum). For example, a sample in the left cortex could be assigned only to an atlas parcel in the left cortex (Arnatkevičiūtė et al., 2019). If a brain region was not assigned a sample from any donor based on the above procedure, the tissue sample closest to the centroid of that parcel was identified independently for each donor. The average of these samples, weighted by the distance between the parcel centroid and the sample, was calculated across donors to estimate the parcellated expression values for the missing region. Tissue samples that were not assigned to a brain region in the provided atlas were discarded.

To address intersubject variation, tissue sample expression values were normalized across genes using a robust sigmoid function (Fulcher, Little, & Jones, 2013):

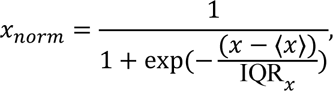

where ⟨*x*⟩ is the median and IQR*_x_* is the normalized interquartile range of the expression of a single tissue ample across genes. Normalized expression values were then rescaled to the unit interval:

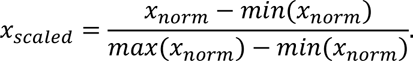

The gene expression values were normalized across tissue samples using the same procedure. Samples assigned to the same brain region were averaged separately for each donor and then across donors. The fully pre-processed gene data included a 426 by 15632 matrix of microarray gene expression data.

#### Identification of genes correlated with dynamic lateralization states

We performed correlation analyses between each group-averaged dynamic lateralization state and the Allen Human Brain spatial gene expression patterns. For each dynamic lateralization state, we calculated the Pearson correlation between each 426-region gene expression vector and the 426-region dynamic lateralization state vector. Multiple comparisons were corrected using BH-FDR corrections (q value < 0.05). Genes that exhibited significant correlations with lateralization states were identified as dynamic lateralization-related genes.

#### Gene set enrichment analysis

To provide further biological insights into the genes that exhibited significant correlations with the lateralization states, we conducted gene set enrichment analysis (GSEA) with FUMA, an integrative web-based platform (K. Watanabe, Taskesen, Van Bochoven, & Posthuma, 2017). We employed the GENE2FUNC function to evaluate the enrichment of the dynamic lateralization-related genes. In the enrichment analysis, hypergeometric tests were performed to determine if the mapped genes were overrepresented in any of the predefined gene sets. Gene sets were obtained from the genes reported in the GWAS Catalog (MacArthur et al., 2017). Multiple test correction was conducted using BH-FDR corrections with an adjusted p-value cutoff of 0.05 and a minimum of 2 overlapping genes.

### Comparison of dynamic lateralization states identified during the resting state and the processing of Chinese natural speech stimuli

The dynamic brain lateralization was analysed using resting-state fMRI data collected from a sample of 991 participants enrolled in the HCP cohort. Detailed algorithm parameters can be found in a previous study (Wu et al., 2022). We quantified the similarity between the group-averaged dynamic lateralization states observed during the Chinese natural listening task and those observed during the resting state.

Furthermore, we statistically analysed the temporal properties of the dynamic lateralization states identified during the resting state, including the occurrence rate and mean dwell time. To investigate potential disparities in these temporal properties between sexes, we employed the following linear mixed-effects model, with age, education level, race, and grey matter volume (GMV) as covariates and the family ID as a random effect:

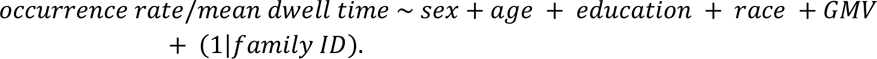

## Acknowledgements

This work was supported by the Natural Science Foundation of China (grant numbers 81901826, 61932008, 62076068, 82271627), the Natural Science Foundation of Shanghai (grant numbers 19ZR1405600, 20ZR1404900), the National Key R&D Program of China (grant numbers 2018YFC1312900 and 2019YFA0709502) the Shanghai Municipal Science and Technology Major Project (grant number 2018SHZDZX01), the ZJ Lab, Shanghai Center for Brain Science and Brain-Inspired Technology and the 111 Project (grant number B18015), and the National Science and Technology Council (Taiwan, grant number NSTC 112-2221-E-033-019).

## Authors’ contributions

All authors had full access to the data in the study and accepted the responsibility to submit it for publication. M.C. and J.F. proposed the study. R.Z. analysed the data. S.G. and W.G. prepared the experimental materials and collected the fMRI data. S.G. preprocessed the data. X.W. and X.Z. assisted in the data analysis. R.Z., M.C., J.F., J.L., X.C., and J.Z. contributed to the interpretation of results. R.Z. and M.C. drafted the manuscript. R.Z. and M.C. edited the manuscript. R.Z. contributed to visualization. All authors approved the manuscript.

## Code availability

Python 3.9.7 was utilized for the analysis of audio texts using natural language processing (NLP) techniques. The NLP techniques applied in this study included the "Jieba" Chinese text segmentation system (https://github.com/fxsjy/jieba/) and text vectorization (https://github.com/Embedding/Chinese-Word-Vectors). Gene expression data were processed in Python using the abagen toolbox (version 0.1.3; https://github.com/rmarkello/abagen). MATLAB 2018b was utilized to calculate the whole-brain dynamic laterality index (DLI) and perform statistical analyses. The MATLAB Brain Connectivity Toolbox (https://sites.google.com/site/bctnet/) was used to perform community detection based on the whole-brain DLI values to identify the dynamic lateralization states and spatial communities of the states. In addition, the MATLAB functions "linkage" and "fitglme" were utilized for the hierarchical clustering of the vectorized articles processed through NLP techniques and for the implementation of the linear mixed-effects models, respectively.

