## Supplementary Figures 1-4 and Supplementary Table 1 for "Dynamics of Brain Lateralization during Chinese Natural Speech Processing under the Influence of Sex Hormones: a 7T fMRI study"

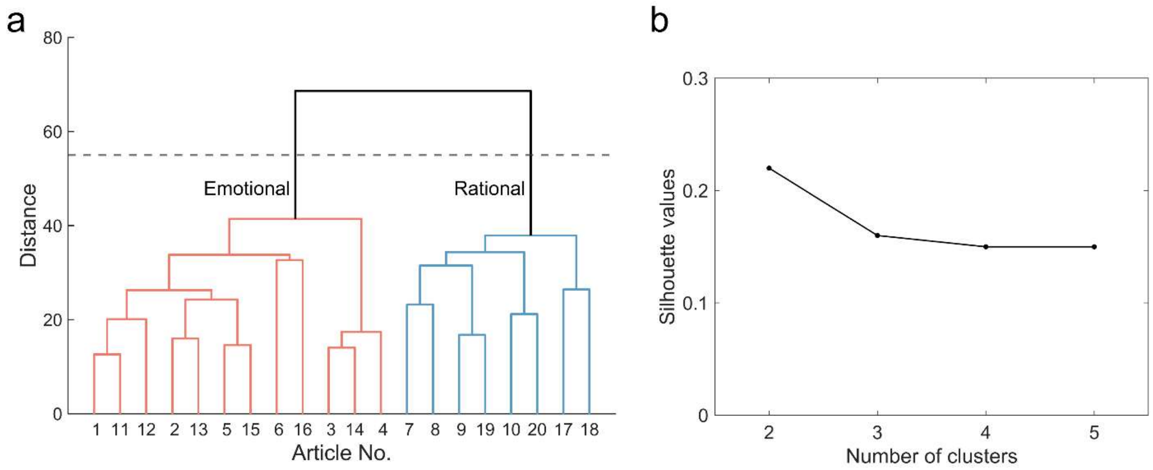


**Figure S1. The hierarchical clustering for article classification.** Two distinct categories were identified. (a) The dendrogram for the clustering results. The two categories of articles were manually labelled as articles with an emotional narrative tone (12 articles) and articles with a rational narrative tone (8 articles). (b**)** An assessment of the optimal number of clusters based on the silhouette criterion. This evaluation confirms the suitability of classifying articles into two categories, as indicated by the highest silhouette value.

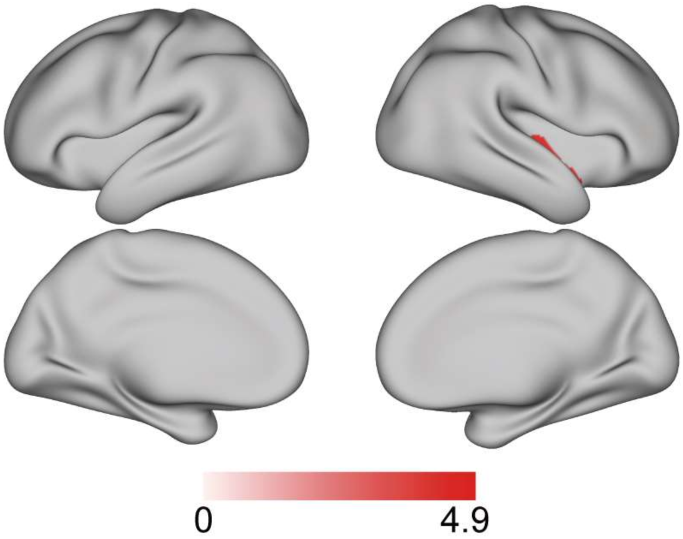


**Figure S2. Differences in the laterality fluctuation between sexes (males > females) during the processing of emotional articles.** Only one region within the right insular cortex (i.e., area PoI1) showed significant sex differences in the LF. The statistical analyses were conducted using linear mixed-effects models, with age and the number of time windows as covariates. The linear mixed-effects model was fitted for the LF of each brain region, using the BH-FDR correction for multiple comparisons (q value < 0.05).

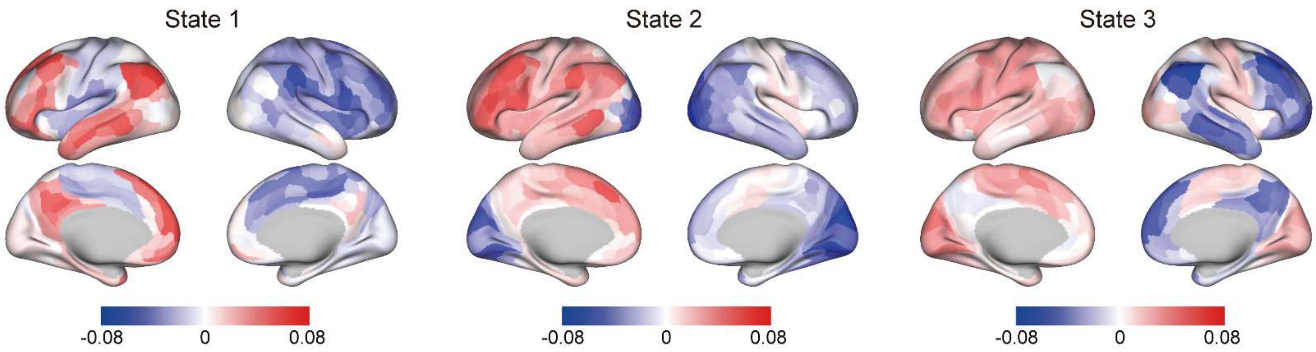


**Figure S3. Three dynamic laterality states identified using resting-state fMRI data with the Human Connectome Project (HCP) cohort.** Among these three states, State 1 and State 3 exhibited significant similarities with State 1 ($r=0.79$, $p=6.4\times10^{-79}$) and State 2 ($r=0.79$, $p=5.3\times10^{-79}$) of our task fMRI findings, respectively, which were identified during the processing of the Chinese natural speech stimuli.

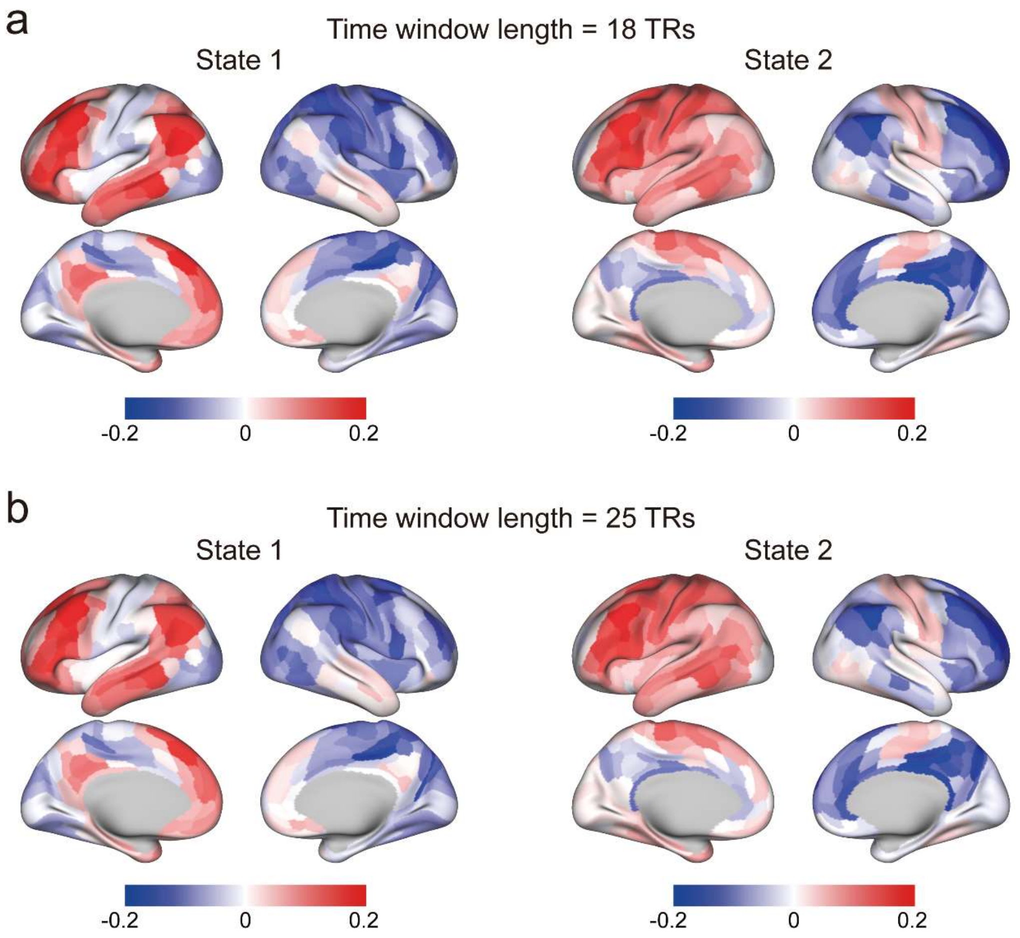


**Figure S4. Validation of dynamic laterality states over varying time window lengths.** Results under both 18 TRs and 25 TRs time window lengths consistently confirm the identification of two distinct dynamic laterality states. (a) The two dynamic laterality states identified using an 18-TR time window. These states exhibit a remarkable similarity to those identified with a 20-TR time window (for State 1, $r=0.99$, $p<1.0\times10^{-15}$; for State 2, $r=0.99$, $p<1.0\times10^{-15}$). (b) The two states identified with a 25-TR time window, closely resembling those identified with a 20-TR time window (for State 1, $r=0.99$, $p<1.0\times10^{-15}$; for State 2, $r=0.99$, $p<1.0\times10^{-15}$).

### Tables

**Table S1 Summary of each article of the Chinese natural speech stimulus.**

| No. | Title | Topic | Synopsis |
|---|---|---|---|
| 1 | The Rabbit Who Went to the Moon | fairy tale | This story narrates the tale of the space agency's quest to find a rabbit to accompany the Brave spacecraft on a journey to the moon, where it would be left behind for scientific experiments. A white rabbit is chosen and undergoes a unique adventure, ultimately returning to Earth as an unexpected hero and global sensation. |
| 2 | Severing the Friendship | fable | This story narrates the close friendship between Guan Ning and Hua Xin during their youth, which is put to the test when they discover gold and encounter a procession of a high-ranking official. Guan Ning emphasizes hard work and moral values, while Hua Xin is tempted by wealth without labor and material luxury. In the end, Guan Ning decides to sever their friendship because of their differing aspirations and values, symbolized by cutting a mat in half. |
| 3 | The Family | novel | The story revolves around Gao Juexin, the elder brother of Gao Juemin. Despite sharing the same mother and household, their circumstances differ greatly. Gao Juexin is the eldest son and the firstborn in the extended family, shaping his destiny from birth. Gifted with good looks and intelligence, he earns admiration from his parents and teachers. However, his future takes an unexpected turn when his mother passes away and his father arranges a marriage for him, shattering his dreams of further education and the love he had for another woman. |
| 4 | The Class Adviser | novel | The story unfolds as Song Baoqi, a former delinquent, is released from detention, leading to his enrollment in Zhang Junshi's class. Despite doubts from his colleagues, Teacher Zhang takes on the challenge of guiding and mentoring Song Baoqi, showcasing his commitment to making a difference in the troubled student's life. This story explores the complexities of education, understanding, and trust. |
| 5 | The Color of Light | essay | When Cézanne painted apples in blue, it altered people's perceptions of color, from Matisse's blue sunflowers to Picasso's vibrant red figures. Artists chase absolute reality, though it is often elusive. The author recalls a moment at a party where the blue light transformed apples, evoking the essence of Cézanne's art. This essay explores how light and atmosphere influence our perception of color and how modern artists challenge traditional notions of reality. |

| | | | |
|---|---|---|---|
| 6 | The Hometown Banyan Trees | essay | The author reflects on two venerable banyan trees near their residence, offering refreshing shade and a haven for children's play. These banyans evoke cherished memories, and the author frequently visits them with their child, reliving moments from the past, including the legend of a snake spirit. These banyan trees symbolize the unity and warmth of the village community, enduring even as time passes and individuals venture far from their hometown. |
| 7 | Police Arrest 26 Suspects After Airing of "Crazy Personal Information Black Market" | news | In February, CCTV News Channel broadcasted a program revealing the illegal trade of personal information online. Following this report, the Public Security Ministry formed a special investigative team. After nearly three months of investigation spanning Beijing and other regions, the case has been successfully resolved, resulting in the apprehension of 26 individuals connected to criminal activities involving the theft and sale of personal information. |
| 8 | Huabei Connects to the Central Bank's Credit System, Gradually Covering All User Groups | news | This news tells that as early spending becomes a daily habit, digital financial products like Huabei, Jiebei, JD Bai Tiao, and Weilidai gain popularity alongside credit cards. Recently, Huabei has upgraded its services by integrating with the central bank's credit system. While not all users are included, specific groups have been integrated, enabling the monthly reporting of their financial information. This step aims to enhance credit assessments' accuracy and improve the overall credit system. |
| 9 | Petroleum Resources | scientific explanatory essay | Petroleum is formed underground under high pressure over extended periods and requires drilling for extraction, followed by separation and processing in refineries. Despite the increasing challenges of petroleum exploration, there are substantial undiscovered petroleum resources. However, petroleum extraction can have environmental impacts, and advanced technology and strict laws are helping to control these adverse impacts on the environment. |
| 10 | Bombed by Negative News, Here's How to Maintain Mental Health | news | When bombarded by negative news, people often feel their mood affected, but this is a natural response of the human brain to threatening information. Over-empathizing or getting too immersed in negative comments can exacerbate negative emotions. Media literacy and the ability to discern false information become crucial, and active participation in public discussions and actions helps alleviate feelings of powerlessness. Focusing on change and taking action, rather than passively succumbing to news, is the best way to mitigate the sense of helplessness brought about by "distant cries." |

| | | | |
|---|---|---|---|
| 11 | School in the Clouds | fairy tale | Laughing Cat, Mouse Qiu Qiu, and the narrator visit Mouse Xiao Bai's hostess's villa, where they are initially alarmed by strange voices. However, they soon realize the villa is empty except for Lory, a voice-mimicking parrot. A sudden storm separates Laughing Cat and Mouse Qiu Qiu, but when Laughing Cat awakens, he reunites with Mouse Xiao Bai and together they rescue Mouse Qiu Qiu from a tree. Little White then shares the enchanting story of how he and his hostess arrived, riding a magical oiled-paper umbrella, and invites Laughing Cat to meet her the next day. |
| 12 | The Battle of the Yellow Emperor Against Chi You | fable | In this ancient Chinese myth, Chi You, a descendant of the Flame Emperor, waged a war against the Yellow Emperor but was captured. With the help of the Kuafu tribe, Chi You's forces regained strength, challenging the Yellow Emperor. Seeking guidance from Xuan Nu, a celestial being, the Yellow Emperor obtained a magical sword and strategic knowledge. Equipped with these newfound resources, the Yellow Emperor reversed the course of the war, defeating Chi You and his allies. Chi You was executed, and his shackles became a grove of enduring red maple trees, signifying his legacy. This story illustrates the importance of wisdom, strategy, and unity in overcoming formidable challenges. |
| 13 | Chronicle of a Blood Merchant | novel | This story narrates how a man named Xu Sanguan leisurely lounges in a melon field, savoring the sweet watermelon and engaging in a conversation about different melon varieties with his uncle. Unexpectedly, Xu Sanguan declares his desire to get married. Shortly after this declaration, he crosses paths with a young woman named Xu Yulan, and a newfound bond blossoms between them. |
| 14 | Ordinary World | novel | The story describes Sun Shaoping's challenging life as a high school student from a poor rural background. He struggles with hunger and self-esteem issues due to his family's financial difficulties, yet he feels a sense of pride for having the opportunity to attend school in a larger world. Despite the hardships, he cherishes the experience of leaving his remote village for a broader horizon. |
| 15 | Poetic Night Rain | essay | This essay reflects the profound impact of night rain on travelers, invoking both longing for home and a sense of tranquility. It illustrates how this natural phenomenon can shape emotions and even alter the course of history, affecting generals, advisors, kings, and heroes. Through the poet's perspective, night rain connects life's everyday experiences to deeper meanings, making it a rich source of inspiration. |

| | | | |
|---|---|---|---|
| 16 | The Five Key Flavors of Chinese Food | essay | This article highlights the diverse palate of people across China. The tastes of Chinese food are traditionally categorized into five flavors: sour, sweet, bitter, spicy, salty, encompassing the vinegar affection of Shanxi residents, the spicy cravings of Sichuan locals, the fondness for sweets among Guangdong inhabitants, and the distinctive flavors of Beijing's stinky tofu, etc. It underscores the richness of Chinese cuisine and the depth of its culinary culture. |
| 17 | Safeguarding the Vital Thread of Ethnic Unity | news | This article showcases how Yunnan, a province rich in ethnic diversity, has made ethnic unity and progress a top priority, in line with the President of China, Xi Jinping's vision. Through focused poverty reduction initiatives, improved infrastructure, and economic diversification, Yunnan has uplifted its ethnic minority communities, bolstered by legal measures to safeguard and advance ethnic development, solidifying its unwavering dedication to ethnic unity. |
| 18 | Chinese President Xi Jinping Empowers Entrepreneurs for Greater Economic Impact | news | Chinese President Xi Jinping chaired a symposium with entrepreneurs, recognizing their vital role in China's economy, especially during the COVID-19 pandemic. He encouraged them to uphold values like innovation, integrity, and social responsibility while addressing domestic and global challenges and promoting economic cooperation through reforms for a thriving global economy. |
| 19 | Flowing Water on Mars | scientific explanatory essay | This article discusses how Mars' photos imply a history of substantial liquid water, seen through river-like runoff and outflow channels. These signs point to a thicker atmosphere and warmer climate about 4 billion years ago. Although debates continue regarding ancient Martian oceans and lakes, the presence of outflow channels stands as compelling evidence of prior water abundance, mostly preserved in subterranean ice, especially within polar regions. |
| 20 | How Does the Brain Perceive the Passage of Time? | scientific explanatory essay | This article delves into the intricate ways our brains perceive time, highlighting the complexity of our internal timing systems. Different brain regions may host various internal clocks operating at different speeds, which collectively enable our ability to process time. Experiments, including neuroimaging and animal studies, indicate that neurons encode relative time rather than absolute time. This understanding may offer insights into broader questions regarding how the brain constructs our perception of reality. |
